# Strategies of *Mycoplasma ovipneumoniae* to evade immune clearance by alveolar macrophages

**DOI:** 10.1101/2021.06.28.450257

**Authors:** Liyang Gao, Kai Zhang, Ying Zhang, Chunji Ma, Xiaoyu Zhou, Min Li

## Abstract

Chronic nonprogressive pneumonia is a prevalent disease that infects many young sheep. *Mycoplasma ovipneumoniae* was isolated from the lungs of sheep with chronic nonprogressive pneumonia. Evidences showed that it might associate with the development and duration of chronic pneumonia, moreover, sheep infected with *M. ovipneumoniae* are easily infected by other organisms, suggesting that *M. ovipneumoniae* may play an immunosuppressive role during infection. However, the mechanism is still poorly understood. The infection occurs in the airway, where resident alveolar macrophages first encounter *M. ovipneumoniae*. Therefore, primary alveolar macrophages (AMs) were collected from the lungs of healthy adult sheep, and the (iTRAQ) protein assay was used to investigate the immunosuppressive effects of *M. ovipneumoniae* on sheep AMs. The RAW264.7 cells were used to confirm the findings. The results showed that *M. ovipneumoniae* promoted higher expression of anti-apoptotic proteins and lower expression of apoptosis-related proteins in the infected AMs. Moreover, the number of infected AMs increased. However, *M. ovipneumoniae* reduced ATP levels in AMs and impaired late endosome maturation and phagolysosome fusion. Furthermore, *M. ovipneumoniae* inhibited the autophagy pathway via the Akt-mTOR axis in AMs. These findings indicated that *M. ovipneumoniae* had distinctive strategies to evade elimination caused by the AMs. The findings might explain the chronic infection and co-infection in sheep infected by *M. ovipneumoniae*.

## 1 INTRODUCTION

Chronic nonprogressive pneumonia is a prevalent disease that infects many young sheep. *Mycoplasma ovipneumoniae (M. ovipneumoniae)*, which has been isolated from the lungs of sheep with chronic nonprogressive pneumonia, is associated with the development and duration of chronic pneumonia. *M. ovipneumoniae* has been found in goats (Livingston & Gauer, 1979; Mohan, Obwolo, & Hill, 1992; Rifatbegovic, Maksimovic, & Hulaj, 2011; Rong et al., 2014), bighorn sheep (Besser et al., 2014), moose, caribou, mule, white-tailed deer, and mule deer (Highland et al., 2018) besides sheep. Previous studies of the pathogenic mechanisms of *M. ovipneumoniae* showed that this bacterium caused direct cell death via the reactive oxygen species (ROS) and the the mitogen-activated protein kinase (MAPK) signaling–mediated mitochondrial apoptotic pathway (Li et al., 2016; Xue et al., 2017). Moreover, the *M. ovipneumoniae* capsular polysaccharide caused the apoptosis of sheep airway epithelial cells via the c-Jun N-terminal kinase (JNK)/P38 MAPK signaling pathway (Jiang et al., 2017; Weiser et al., 2012). Moreover, sheep infected with *M. ovipneumoniae* were easily infected by other organisms, suggesting. that *M. ovipneumoniae* might play an immunosuppressive role during infection in addition to causing airway cell apoptosis.

The airway epithelium functions as a first-line physical barrier during *M. ovipneumoniae* infection. However, alveolar macrophages (AMs), which are innate immune cells, are the host’s first line of defense against pathogens. AMs are tissue-resident phagocytes in the airway lumen and play important roles in regulating airway immunity and conditions, including removing pathogens, producing antibacterial substances, expressing inflammatory factors, and rebuilding the airway environment. Macrophages use two mechanisms to kill pathogens: (1) the pathogens are engulfed via phagocytosis and then degraded by fusing with lysosomes (Sanjuan & Green, 2008); and (2) the pathogens, which invade cells, are degraded via selective autophagy or xenophagy (Levine, Mizushima, & Virgin, 2011). AMs use two important pathways to degrade pathogens: phagocytosis and autophagy. However, microbes have developed strategies to escape degradation by AMs, by either killing AMs (Bewley et al., 2014; Du et al., 2018b; Mehto et al., 2015) or disrupting phagolysosomal maturation and antigen presentation (Holloway, Fleming, & Coulson, 2018; Kissing et al., 2015). A previous study on RAW264.7 macrophages revealed that autophagy was suppressed during *M. ovipneumoniae* infection, but the mechanisms underlying this suppression remain unclear. This study focused on the effects of *M. ovipneumoniae* on sheep primary AMs and confirmed results using RAW264.7 cells.

## 2. MATERIALS AND METHODS

### 2.1 Isolation of primary sheep AMs

Primary AMs were isolated from the lungs of healthy adult sheep brought from a legal sheep slaughtering field. The sheep slaughtering field followed the relevant provisions of the “People’s Republic of China animal epidemic prevention law” and the national declaration of quarantine by the animal health supervision institution. Sterile phosphate-buffered saline (PBS, BIOIND, Shanghai, China) was infused into the trachea. The sheep lungs were serially washed twice, 10 min each time, with 1 L of sterile PBS, and then the lavage fluid was collected. The cell pellets were obtained via centrifugation at 250*g* for 5 min after treatment with red cell lysing reagent (Solarbio Life Sciences, Shanghai, China) for 10 min at 4°C and washing with sterile PBS (Hyclone, Shanghai, China). The cells were suspended in prewarmed Dulbecco’s modified Eagle’s medium (DMEM; Gibco, CA, USA) with 15% fetal bovine serum (Gibco, New Zealand) and 1% penicillin/streptomycin (Hyclone, UT, USA). They were then cultured at 37°C with 5% CO_2_. Next, the semi-attached cells were collected 2 h after plating and then processed with cell characterization.

### 2.2 *M. ovipneumoniae* cultures

The Y98 standard *M. ovipneumoniae* strain was obtained from the National Institutes for Food and Drug Control. Mycoplasma broth base (Hope Bio Technology, Qingdao, China) and fetal equine serum (Solarbio, Shanghai, China) were used to culture *M. ovipneumoniae*. We mixed one volume of *M. ovipneumoniae* with five volumes of fresh mycoplasma broth to thaw *M. ovipneumoniae*. Then it was incubated in an incubator at 37°C (Sanyo, Japan). The *M. ovipneumoniae* population was examined using the color change unit (CCU) assay, which was based on the metabolic activity of *M. ovipneumoniae* cells in the medium. The *M. ovipneumoniae* concentration was the highest when the medium turned yellow (pH <6.8) with phenol red staining (Purcell, Taylor-Robinson, Wong, & Chanock, 1966). The CCU assay was performed with tubes containing a serially diluted mycoplasma suspension (10^1^, 10^2^, 10^3^, 10^4^, 10^5^, 10^6^, 10^7^, and 10^8^), in which the color change was equivalent to 0.5 pH units.

### 2.4 Establishment of *M. ovipneumoniae* infection model *in vitro*

Both primary sheep AMs and Raw264.7 cell line were used to establish *M. ovipneumoniae* infection model. Both primary sheep lung AMs and Raw264.7 mouse macrophages were cultured under the same conditions. Then, *M. ovipneumoniae* strain Y98 (10^7^ CCU/mL, Fig. 1A) was distributed onto dishes containing primary sheep AMs and Raw264.7 mouse macrophages, and the ratio of macrophages and *M. ovipneumoniae* was adjusted to 1:100. *M. ovipneumoniae* was stained with green fluorescence, lipophilic carbocyanine DiOC_18_(3) (Beyotime Biotechnology, China) to monitor the interaction between *M. ovipneumoniae* and macrophages, which could be tracked under fluorescent microscopy. Then, DiOC_18_(3)-stained *M. ovipneumoniae* was washed with PBS and centrifuged at 12,000 rpm for 10 min. This process was repeated three times until no residual DiOC_18_(3) remained in the medium. The labeled *M. ovipneumoniae* was added into the cell culture medium and co-cultured with macrophages for 24 h. Finally, the cover slips were washed with PBS and fixed with 4% paraformaldehyde (Beyotime Biotechnology), prior to observation under a fluorescence microscope. Bafilomycin A (Beyotime Biotechnology) and rapamycin (Beyotime Biotechnology) were added to the cell culture medium to inhibit and stimulate autophagy, respectively.

### 2.3 Protein assay

Proteins were extracted from control AMs and *M. ovipneumoniae*–infected AMs with three biological replicates using the cell lysis buffer for Western blot analysis (Beyotime Biotechnology). The protein quality was tested using a bicinchoninic acid (BCA) protein assay kit (Beyotime Biotechnology). Equal amounts of 5 μg protein from six samples were used. iTRAQ labeling was performed using iTRAQ Reagents-8plex (AB Sciex Inc., MA, USA; Sciex iChemistry, Product Number 4390812) following the manufacturer’s protocol. The samples were digested and labeled with iTRAQ labels. The control samples were labeled with iTRAQ8-113, iTRAQ8-116, and iTRAQ8-117, and the infected samples were labeled with iTRAQ8-118, iTRAQ8-119, and iTRAQ8-121. Two-stage mass spectrometry (MS/MS) was used to label and segregate protein fragments into the reporter, peptide-reactive, and balanced groups. The peptides were incubated with isobaric tags for 2 h at room temperature. Agilent 1260 Infinity high-performance liquid chromatography (Agilent Technologies, CA, USA) was used for strong cationic exchange chromatography. The DIONEX NANO HPLC SYSTEM (ULTIMATE 3000, Thermo Scientific, CA, US) and a Q Exactive mass spectrometer (Thermo Scientific, CA, USA) were used for LC-MS/MS analysis. Then, 1-μg samples were loaded into the DIONEX nano-UPLC system to separate the peptides. The Orbitrap mass analyzer (Thermo Fisher Scientific, CA, USA) was used at a mass resolution of 70,000 FWHM at 400 *m*/*z* in a Q Exactive with full scans (350–1600 *m*/*z*) to perform the mass spectrometry analysis. The LC-MS/MS results were searched against the *Ovis aries* genome protein database of the National Center for Biotechnology Information using Proteome Discoverer software 1.4 (Thermo Scientific). Low-confidence peptides with a global false discovery rate of ≥1% were removed in the protein analysis. Proteins containing at least two unique peptides were quantified. The Gene Ontology (GO) terms were mapped using the GO database (http://www.geneontology.org). Pathway enrichment was analyzed using the Kyoto Encyclopedia of Genes and Genomes database (http://www.genome.jp/kegg/). Some proteins were examined in this study, including B-cell lymphoma 2 (Bcl-2), Bcl-2 homologous antago-nist/killer (Bak), Bcl-2-associated X proten (Bax), Bcl2 Antagonist of Cell Death (Bad), microtubule-associated protein 1 light chain 3 (LC3), autophagy related 5 homolog (Atg5), early endosome antigen 1 (Eea1), NF-κB, 5’-AMP-activated protein kinase (AMPK), c-Jun N-terminal kinase (JUN), AKT serine/threonine kinase 1 (AKT), mechanistic target of rapamycin kinase (MTOR). For Western blot analysis, the following antibodies were used: anti-Bcl-2 (Beyotime; dilution1:500-1:1000), anti-Bak (Thermofisher; dilution1:1000), anti-BAD (Thermofisher; dilution 1:1000), anti-caspase 9 (Proteintech; dilution 1:300-1:1000), anti-Beclin 1 (Proteintech; dilution 1:500-1:2000), anti-LC3B (Beyotime; dilution 1:1000), anti-Atg5 (Proteintech; dilution 1:500-1:2000), anti-phospho-Akt (Proteintech; dilution 1:2000-1:10000), anti-phospho-mTOR (Thermofisher; dilution 1:1000), anti-EEA1 (Proteintech; dilution 1:2000), anti-LAMP1 (Proteintech; dilution 1:2000), anti-LAMP2 (Proteintech; dilution 1:2000), anti-ATP6V1A (Proteintech; dilution 1:2000), and anti-GAPDH (Proteintech; dilution 1:40000). The membrane was incubated with peroxidase-conjugated secondary antibodies, such as HRP Goat anti-Mouse IgG (H+L) (Proteintech; dilution 1:20000), HRP Goat anti-Rabbit IgG (H+L) (Proteintech; dilution 1:20000). ECL Western blot substrate was bought from Thermofisher Scientific.

### 2.4 Immunocytochemistry

Primary sheep AMs and RAW264.7 cells were plated in 24-well plates containing sterile coverslips. AMs were infected with *M. ovipneumoniae* when 70% of the cells were attached to the coverslips. All coverslips were then collected from the plates and fixed with 4% paraformaldehyde for 30 min in room temperature (RT). Thereafter, the cells were permeabilized with 0.1% Triton X-100 (Sigma, 9002-93-1, Shanghai, China) for 15 min in RT and then blocked with 5% goat serum in PBS for 30 min. The primary antibodies were as follows: anti-Bcl-2 (Beyotime; dilution 1:200), anti-BAK (Thermofisher; dilution 1:50), anti-Bax (Thermofisher; dilution 1:50), anti-BAD (Thermofisher; dilution 1:50), anti-caspase 9 (Proteintech; dilution 1:100), anti-Beclin 1 (Proteintech; dilution 1:50-1:500), anti-LC3 (Beyotime; dilution 1:400), anti-Atg5 (Proteintech; dilution 1:50-1:500), anti-phosphp-mTOR (Thermofisher Invitrogen; dilution 1:10-1:50), anti-phospho-NF-Kb (Beyotime; dilution 1:100), anti-phospho-c-Jun(Ser 73) (Thermofisher; dilution 1:100), anti-LAMP1 (Proteintech; dilution 1:200), and anti-LAMP2 (Proteintech; dilution 1:200). The secondary anti-body was anti-Mouse IgG (H+L) Alexa Fluor 488 (Invitrogen; dilution 1:1000), Goat anti-Rabbit IgG (H+L) secondary antibody DyLight 650 (Invitrogen; dilution 1:2000).

### 2.5 Gene expression assay

Total RNA was extracted using a E.Z.N.A. Total RNA extraction kit (Omega, USA) following the manufacturer’s protocols. The purity and concentration of the mRNA samples were examined using a NanoDrop 2000 spectrometer (Thermo Fisher Scientific, DE, USA) and 2% agarose electrophoresis. Reverse transcription was performed using PrimerScript RT Master Mix (TaKaRa, China), and quantitative reverse transcription-polymerase china reaction (RT-qPCR) was performed using SYBR Premix Ex Taq (TaKaRa, China). All procedures were performed following the manufacturer’s protocols.

### 2.6 Cell viability and cell cycle assay

For the cell viability assay, the macrophages were seeded in 96-well plates at a concentration of 1 × 10^4^ cell per well 1 day before the assay. After the treatment, a cell counting kit 8 (CCK-8; Beyotime Biotechnology) was added to 96-well plates incubated at 37 ℃ for 1–4 h, followed by reading under 450 nm in EnSpire microtiter plate spectrophotometer (PerkinElmer, German). A CellTiter-Lumi Luminescent Cell Viability Assay Kit (Beyotime Biotechnology), which showed the ATP level of cells, was also used to examine the cell viability. After the treatment, 100 μL of the reagent was added to 96-well plates, which contained cells and 100 μL of the medium, and then the luminescence was checked using EnSpire microtiter plate spectrophotometer (PerkinElmer, German).

### 2.7 Bacterial killing assay

The macrophages were derived into two groups: *M. ovipneumoniae*–infected group and noninfected group. They were infected with *M. ovipneumoniae* (Y98) for 24 h at 37℃. Then, 10 μg/mL gentamicin (Gibco, USA) was added to the cell culture medium to remove external *Escherichia coli*, and then both noninfected and infected cells were counted and re-plated in a six-well-plate at a density of 5 × 10^5^ cell/mL. *E. coli* DH5α (TransGen Biotech, China) was used to infect both groups for 15 min at 37℃. After 15-min incubation, 50 μg/mL kanamycin (Sigma) was added to the medium to remove the extracellular *E. coli*. The macrophages were lysed with 0.1% (*v*/*v*) Triton X-100 for 15 min and 6 h, respectively, to examine the phagocytosis rate and immune clearance rate. The bacteria inside the macrophages were released and collected by centrifugation at 8000 rpm for 5 min. Finally, the diluted lysate was plated on agar plates overnight, and the colonies were counted.

The bacterial killing was determined using the following formula:
`E. coli survival rate = (Numbers of colony after 6h)/(Numbers of colony after 15min)`X100%

### 2.8 Phagocytosis assay

Immunoglobulin (IgG)-coated magnetic beads (Shengon Biotech, China) and chicken erythrocytes (Solarbio, China) were used to check the phagocytosis of macrophages. The cells were grown to semi-confluence on a six-well plate, and then the medium was replaced with IgG-coated magnetic beads containing serum-free DMEM. Then, the plate was centrifuged at 300*g* for 1 min to allow the precipitation of beads, and then incubated at 37℃ in the presence of 5% CO_2_. After 15 min, excess beads were washed away with PBS three times, and the sample was monitored under the phase-contrast microscope. To monitor the chicken erythrocytes in sheep primary AMs, hematoxylin-eosin (HE) staining was used.

## 3 RESULTS

### 3.1 *M. ovipneumoniae* infection model of primary sheep AMs and Raw264.7 cells

Primary sheep lung AMs were isolated from infused airways. After 2-h culture, only semi-attached cells were collected to process with morphological analysis, immunocytochemistry, and function assay. The images showed that all cells were round in shape, and up to 90% of the primary sheep AMs were CD14-positive cells (Supplementary Fig. 1A). Moreover, the chicken erythrocytes were used to confirm the phagocytosis of primary sheep lung AMs. HE staining images showed that the primary sheep AMs engulfed the chicken erythrocytes (Supplementary Fig. 1B).

*M. ovipneumoniae* was stained with DiOC_18_(3) and co-cultured with primary sheep AMs and Raw264.7 for 24 h to check the interaction between *M. ovipneumoniae* and macrophages. The images showed that the macrophages engulfed *M. ovipneumoniae*, storing them inside the cytoplasm (Fig. 1).

### 3.2 *M. ovipneumoniae* infection reduced the apoptosis of macrophages

The cell viability of the primary AMs was checked after *M. ovipneumoniae* infection for 12 and 24 h. The result showed that *M. ovipneumoniae* infection did not reduce the cell population of the primary AMs, but slightly increased the cell number of primary AMs (Fig. 2A). The expression of key anti-apoptotic factor Bcl-2 significantly increased after infection (Fig. 2B), while the expression of pro-apoptotic genes *BID* and *BAD* decreased (Fig. 2C). RAW264.7 cells were infected for 24 h to further confirm the anti-apoptotic effects of *M. ovipneumoniae* on macrophages. The cell population increased after *M. ovipneumoniae* infection(Fig. 3A); however, the ATP level decreased significantly (Fig. 3B). In addition, the cell cycle of RAW264.7 showed no obvious changes after infection (Fig. 3C-D). Furthermore, the levels of pro-apoptotic factors decreased in the *M. ovipneumoniae*–infected groups compared with the non-infected group (Fig.3E).

### 3.3 *M. ovipneumoniae* evaded the intercellular pathogen clearance of macrophages by reducing autophagy via the Akt-mTOR signaling pathway

The density of *M. ovipneumoniae* was determined 24 h after infection to examine the effect of primary sheep AMs on the clearance of *M. ovipneumoniae* during infection. In this study, two chemicals were used, bafilomycin A, which was used as an autophagy pathway inhibitor, and rapamycin, which served as an autophagy stimulator. Bafilomycin A and rapamycin were used to treat infected primary sheep AMs. Then, *M. ovipneumoniae* was released from primary sheep AMs using a high-speed centrifuge (12,000 rpm, 10 min). The cytoplasmic *M. ovipneumoniae* was cultured at 37℃ for a week until the medium of some vials turned yellow. *M. ovipneumoniae*–infected primary AMs treated with rapamycin had the lowest cytoplasmic *M. ovipneumoniae* density (1X10^5^ CCU/mL); however, the non-treated groups and bafilomycin A-treated groups shared the similar density (1X10^6^ CCU/mL) of cytoplasmic *M. ovipneumoniae* (Fig. 4A-B).

The suppressive effect of cytoplasmic *M. ovipneumoniae* on the autophagy of primary sheep AMs during pathogen clearance was explored. Then, the expression of autophagy-related proteins in primary sheep AMs, including Beclin 1 and LC3, was examined. The levels of these proteins increased during the initial infection (6–24 h) but decreased after 36 h (Fig. 4C-D).

The bacterial killing assay was performed using Raw264.7 cells to confirm the reducing effect of *M. ovipneumoniae* on the pathogen clearance of macrophages (Fig.5A, Supplementary Fig.2). The results showed that *M. ovipneumoniae* infected primary sheep AMs, whose phagocytosis significantly increased after co-culturing with *E. coli* for 15 min (Fig. 5B). The *E. coli* survival rate in *M. ovipneumoniae* infected macrophages was higher than that in the non-infected macrophages. However, rapamycin could reduce the survival rate of *E. coli* in *M. ovipneumoniae*–infected macrophages (Fig.5C).

The expression of autophagy-related proteins in Raw264.7 cells was examined to further explore the molecular mechanism of *M. ovipneumoniae*–induced autophagy suppression in macrophages. First, the expression of Atg5, LC3, and Beclin 1 increased, but the expression of p-Akt and p-mTOR decreased during the early infection (24 h). Then, *M. ovipneumoniae* was withdrawn from the medium, followed by the induction of autophagy via serum deprivation. Both the noninfected and infected Raw264.7 cells were continually cultured for 24 h (Fig. 6A). In contrast, the levels of p-Akt and p-mTOR increased with time, and the expression of Atg5, Lc3II, and Beclin 1 decreased after 48 h (Fig. 6B).

### 3.5 *M. ovipneumoniae* impaired phagocytic clearance by damaging phagolysosomal maturation in macrophages

IgG-coated beads (500 nm) were co-cultured with *M. ovipneumoniae*–infected RAW264.7 cells for 15 min to examine the phagocytosis of infected cells. The images showed that although the morphology of infected RAW264.7 changed, these cells could engulf beads, indicating that the infected macrophages still had phagocytic function (Supplementary Fig.3).

Next, iTRAQ labeling was used to examine the protein expression in primary sheep AMs. The clustering of differential protein expression is shown in Supplementary Fig.4. The expression of lysosomal membrane proteins decreased (Fig. 7A), including major lysosomal membrane protein 2 (SCARB2), minor lysosomal membrane protein Niemann-pick disease type C (NPC) intracellular cholesterol transporter 2, and ceroid-lipofuscinosis neuronal protein 5 (CLN5). Moreover, the levels of lysosome acid hydrolases, including protease cathepsins (CTSZ, CTSS, and CTSK), glycosidase lysosomal alpha-glucosidase (GAA), sulfatase N-acetylgalactosamine-6-sulfatase (GALNS), lipase lysosomal acid lipase ester hydrolase precursor (LIPA), and acid ceramidase (ASAH1), decreased (Fig. 7B). Interestingly, the levels of mannose-6-phosphate (M6P) receptor on the Golgi apparatus, which delivered lysosomal enzymes to the lysosomes, decreased after infection. The levels of H^+^-ATPase (ATP6V1A) on the membranes of early and mature phagosomes and phagolysosomes also decreased after *M. ovipneumoniae* infection (Fig. 7C). The levels of early endosome antigen 1 (EEA1) and beta-tubulin (TUBB) increased (Fig. 7C). The levels of phagocytosis-promoting receptors, including integrin alpha-M precursor (ITGAM), thrombospondin 1 (THBS1), and mannose-binding protein A (MBLA), increased in AMs after *M. ovipneumoniae* infection (Fig. 7C). RT-qPCR was performed to confirm the results of iTRAQ proteomic analysis. The mRNA expression level followed a similar pattern as the protein expression level (Supplementary Fig.5).

Then, the expression of Eea1, Lamp1, and Lamp2 was examined in Raw264.7 cells. The Western blot analysis results showed that Eea1 expression increased, suggesting that the *M. ovipneumoniae* infection did not influence early endosome formation. In contrast, Lamp1 and Lamp2 protein expression decreased in *M. ovipneumoniae*–infected cells, suggesting that *M. ovipneumoniae* infection might damage late endosome/lysosome formation (Fig. 8A-B). The levels of H^+^-ATPase normally found on cytolysosome and autolysosome membranes significantly decreased (Fig. 8A-B). The ICC images also showed that the levels of both Lamp1 and Lamp2 decreased in *M. ovipneumoniae*–infected RAW264.7 cells (Fig. 8C-D). Moreover, the *M. ovipneumoniae*–infected cells showed large vesicles inside the cytoplasm, suggesting that *M. ovipneumoniae* might store inside late phagosomes (Fig. 8E).

## 4 DISCUSSION

### 4.1 *M. ovipneumoniae* hid in the cytoplasm of macrophages to avoid elimination

The threats of pathogenic microorganisms to immune system are different. Some pathogens cause inflammation via stimulating innate immune cells, and some pathogens cause apoptosis of immune cells. For example, *Leptospira interrogans* causes the apoptosis of macrophages by triggering DNA fragmentation (Hu et al., 2017). *Helicobacter pylori* also causes apoptosis in macrophages by secreting protein HP1286 (Tavares & Pathak, 2017). *Mycobacterium avium* induces macrophage apoptosis by activating JNK and enhancing ROS production (Lee et al., 2016). *Mycoplasma bovis* increases slight cytotoxicity and apoptosis in macrophages (Bürgi et al., 2018). The lipopolysaccharide from *Leptospira interrogans* also causes macrophage apoptosis (Du et al., 2018a). *M. ovipneumoniae* caused apoptosis in sheep airway epithelial cells via ROS-induced pathways and the ERK-mediated mitochondrial pathway (Jiang et al., 2017; Li et al., 2016; Xue et al., 2017).

However, different from other respiratory tract pathogens such as *Mycobacterium tuberculosis*, *M. ovipneumoniae* infection did not cause apoptosis or necrosis of macrophages. Instead, *M. ovipneumoniae* infection increased the AM population, which was not due to cell cycle changes but due to the inhibition of apoptosis. In our study, we found that *M. ovipneumoniae* infection decreased expression of apoptosis-related proteins, while the anti-apoptosis pathway was stimulated in both primary sheep lung AMs and RAW264.7 cells. Moreover, we found *M. ovipneumoniae* could survival inside the AM cytoplasm, suggesting that *M. ovipneumoniae* might not suppress the immune response via direct cell death of AMs. Instead of direct killing, *M. ovipneumoniae* might hide inside the macrophages. This survival strategy was similar to that of intracellular pathogenic microbes such as *Chlamydia trachomatis* and *Edwardsiella tarda* (Al-Zeer et al., 2017; Nolan et al., 2016). Moreover, our previous study showed that *M. ovipneumoniae* could invade to alveolar epithelial cells (GAO et al., 2020), suggest that *M. ovipneumoniae* shares some characters of Intracellular bacteria, which have not been reported in detail.

### 4.2 *M. ovipneumoniae* inhibited autophagy via the Akt/mTOR signaling pathway in macrophages

Autophagy is another lysosomal degradation pathway in cells. Even if *M. ovipneumoniae* hiding inside the cytoplasma of AMs, the AMs can still degrade it via autophagy. However, our study showed that *M. ovipneumoniae* might inhibit autophagy of AMs. Results showed the expression of autophagy -related proteins involved in autophagosomal membrane initiation, elongation, and formation decreased after infection. Moreover, we found that Calreticulin decreased in infected AMs. Calreticulin is a conserved calcium-binding protein normally used as an the endoplasmic reticulum (ER) marker (Liu, Xu, & Zhang, 2013), and ER is a major membrane source of autophagosomes (Bento et al., 2016). The autophagy could be inhibited via calreticulin knockout (Chao et al., 2010; Feng, Chen, Weissman-tsukamoto, Volkmer, & Yi, 2015). Therefore, calreticulin on the cell surface is important for macrophage phagocytosis. These evidences suggested that *M. ovipneumoniae* might inhibit phagocytosis and autophagy by decreasing expression of calreticulin.

We also found that LC3 expression decreased after *M. ovipneumoniae* infection, suggesting that LC3 might a target of *M. ovipneumoniae*. LC3 is involved in both phagocytosis and autophagy, and LC3II is the membrane-bound form of LC3. Phagocytosis must recruit LC3II to single-membrane phagosomes (Martinez et al., 2011), and LC3II must bind with the canonical autophagosomal double membrane during autophagy (Pimentel-Muiños & Boada-Romero, 2014). Some pathogens interrupt LC3 recruitment to the membrane compartments, for example, *Legionella pneumophila* evades host autophagy using the RavZ protein N-terminus to recognize membrane LC3 and then deconjugates it on autophagosomal membranes (Yang, Pantoom, & Wu, 2017). Our finding showd that *M. ovipneumoniae* might suppress the autophagy pathway via reducing LC3.

The Akt-mTOR pathway is important in regulating autophagy, and rapamycin is widely used to inhibit the Akt-mTOR pathway via binding to mammalian target of rapamycin complex 1(mTORC1). We found that intracellular *M. ovipneumoniae* damaged the autophagy pathway via the Akt-mTOR axis. Instead, stimulating autophagy by rapamycin reduced the *M. ovipneumoniae* population in AMs. Some viruses have developed strategies to inactivate the autophagy pathway via the Akt-mTOR axis (B. Hu et al., 2015; Huang et al., 2013). These evidences suggested that cytoplasmic *M. ovipneumoniae* might reduce the clearance by inhibiting the autophagy pathway.

### 4.3 *M. ovipneumoniae* impaired the phagocytosis of macrophages

In this study, both iTRAQ labeling and RT-qPCR were used to monitor the protein and mRNA expression in sheep primary AMs and Raw264.7 macropahges after *M. ov*i*pneumoniae* infection, respectively. We found that phagocytosis of macropahges were damaged by *M. ov*i*pneumoniae.* Previous studies reported that the microtubule cytoskeleton played an important role during endocytosis in macrophages, and silica particle uptake showed a microtubule-dependent pattern (Gilberti & Knecht, 2014). And the microtubule cytoskeleton is required for non-Fc receptors (FcR)-mediated uptake and GTPase and phosphatidylinositol 3 kinase (PI3K) activation (Yu-shin Sou, Satoshi Waguri, Jun-ichi Iwata, Takashi Ueno, Tsutomu Fujimura et al., 2008). We found that Beta-tubulin expression increased after infection, suggesting that beta-tubulin might be involved in *M. ovipneumoniae* uptake.

*M. ovipneumoniae* uptake might occur through both FcR-mediated and non-FcR-mediated pathways in primary sheep AMs. We found that GTP-binding protein ADP-ribosylation factor 6 (ARF6), which is crucial for membrane recycling during phagocytosis and can be activated during FcR-mediated uptake (Niedergang, Colucci-guyon, Dubois, Raposo, & Chavrier, 2002), was decreased in macrophages after *ovipneumoniae* infection. However, phagocytosis-promoting receptors increased after infection, suggesting that *M. ov*i*pneumoniae* did not impair the early stage of endocytosis.

Nevertheless, *M. ovipneumoniae* altered macrophage function by impairing late endosome maturation and reducing H^+^-ATPase levels on the phagolysosomes. In the next step, the late phagosome fused with the lysosomes via LAMP1 and LAMP2. However, LAMP1 and LAMP2 expression was suppressed in *M. ovipneumoniae*–infected macrophages, indicating that *M. ovipneumoniae* infection impaired phagolysosome maturation. Moreover, we found expression of some lysosomal membrane proteins, including SCARB2, NPC, and CLN5, were significant decreased in *M. ovipneumoniae*–infected macrophages. This result suggesting that *M. ovipneumoniae* infection might impair lysosome function.

Lysosome fusion is the final destination of phagocytosis and autophagy. The late phagosomes and autophagy require a more acidic *pH*. However, the level of vacuolar H^+^-ATPase (v-ATPase) complex, encoded by the *Ovis aries ATP6V1A* gene, significantly decreased in *M. ovipneumoniae*–infected AMs. The H^+^-ATPase complex is critical to pumping protons into lysosomes and acidifying them. The protein assay showed that the v-ATPase subunit decreased bafilomycin-A1, which inhibited autolysosomal formation by blocking the proton transport of v-ATPase, thus maintaining *M. ovipneumoniae* survival in AMs. This finding suggested that *M. ovipneumoniae* altered macrophage functions by impairing late endosome maturation and reducing the H^+^-ATPase on the phagolysosomes.

As a conclusion, we found that although *M. ovipneumoniae* did not reduce the endocytosis, it could inhibit the maturation of late endosome, moreover, it could reduce the function of lysosome by reduce lysosomal membrane proteins and v-ATPase. These findings might explain how *M. ovipneumoniae* suppressed the immune responses of hosts, causing chronic infection and co-infection. These effects also occurred with other microbial infections, for example, the co-infection of *M. tuberculosis* and human immunodeficiency virus (HIV) inhibited AM apoptosis via the Toll-like receptor 2 (TLR2) pathway, enabling these pathogens to establish persistent infections (Mehto et al., 2015).

## 5 CONCLUSION

This study found that *M. ovipneumoniae* escaped the innate immune response using several strategies. First, *M. ovipneumoniae* remained inside AMs and inhibited AM apoptosis. Second, intracellular *M. ovipneumoniae* impaired AM function via different mechanisms, such as decreasing cellular ATP levels, reducing endodermal membrane proteins, arresting phagolysosomal maturation, reducing the H^+^ concentration in the lysosomes, reducing the expression of some lysosomal acid hydrolytic enzymes, and inhibiting AM autophagy through the Akt-mTOR axis (Fig. 9). These findings might explain how *M. ovipneumoniae* escaped the innate immune response and caused chronic infection and co-infections in hosts.

## Abbreviations

AMs: alveolar macrophages
*M.*: *ovipneumoniae*
*Mycoplasma*: *ovipneumoniae*

## 6 ACKNOWLEDGEMENTS

This research was supported by funds from Natural Science Foundation (Grant No.31460039; 31960036), National Chunhui Program of Ministry of Education of China (Grant No.201801).

---

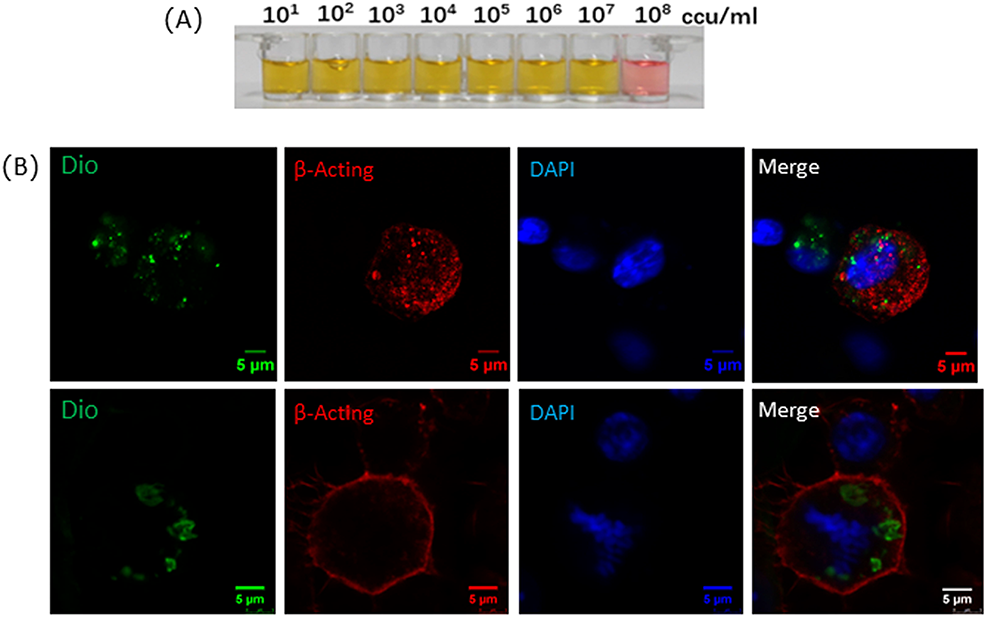

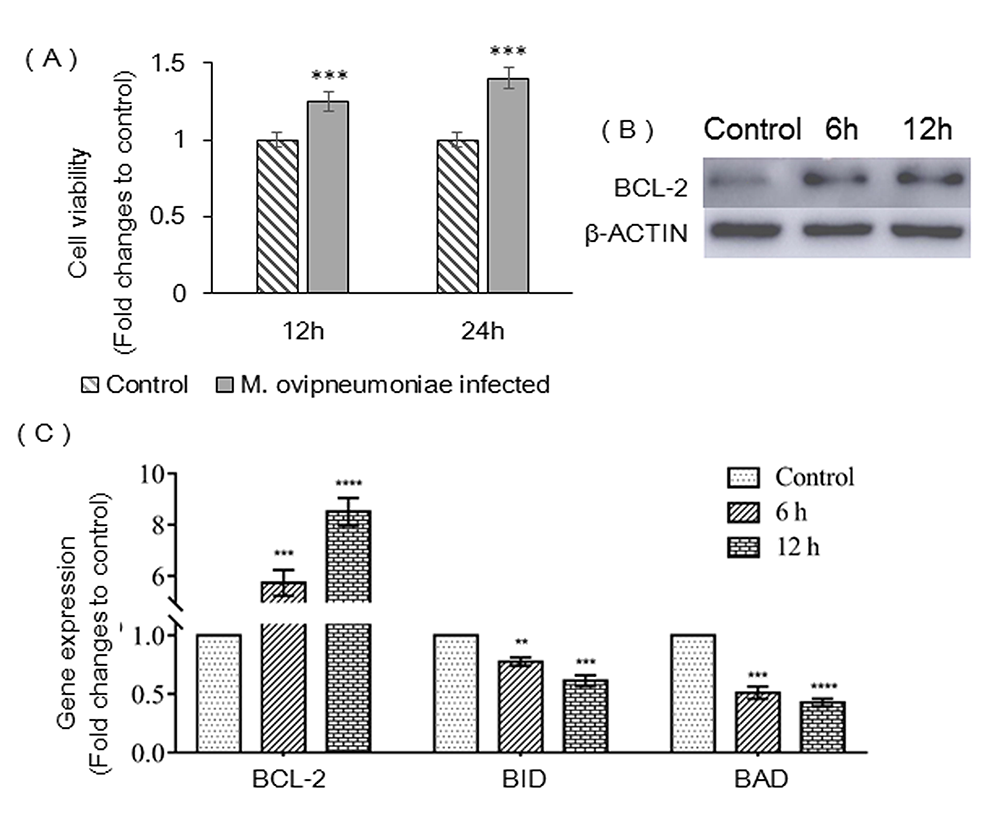

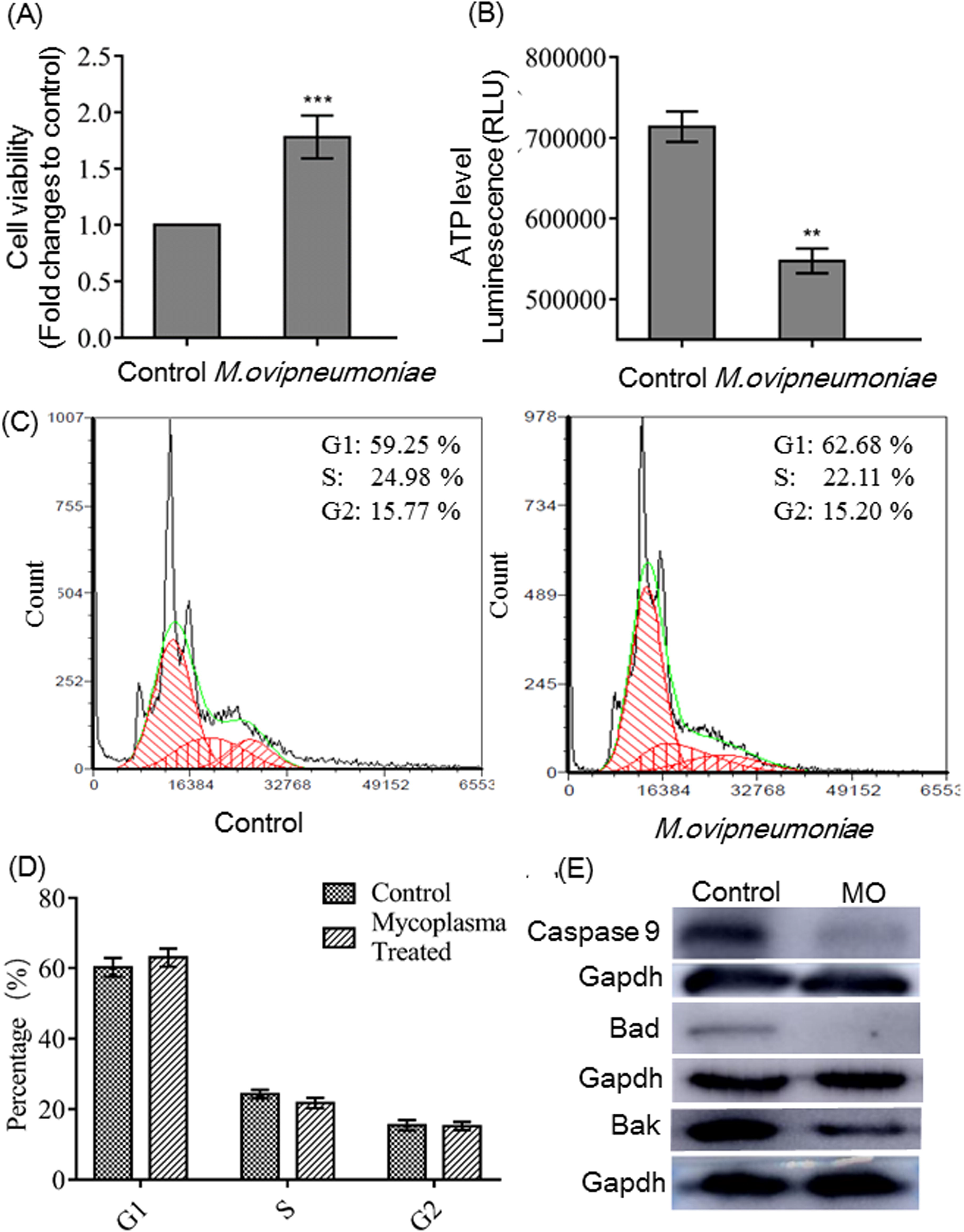

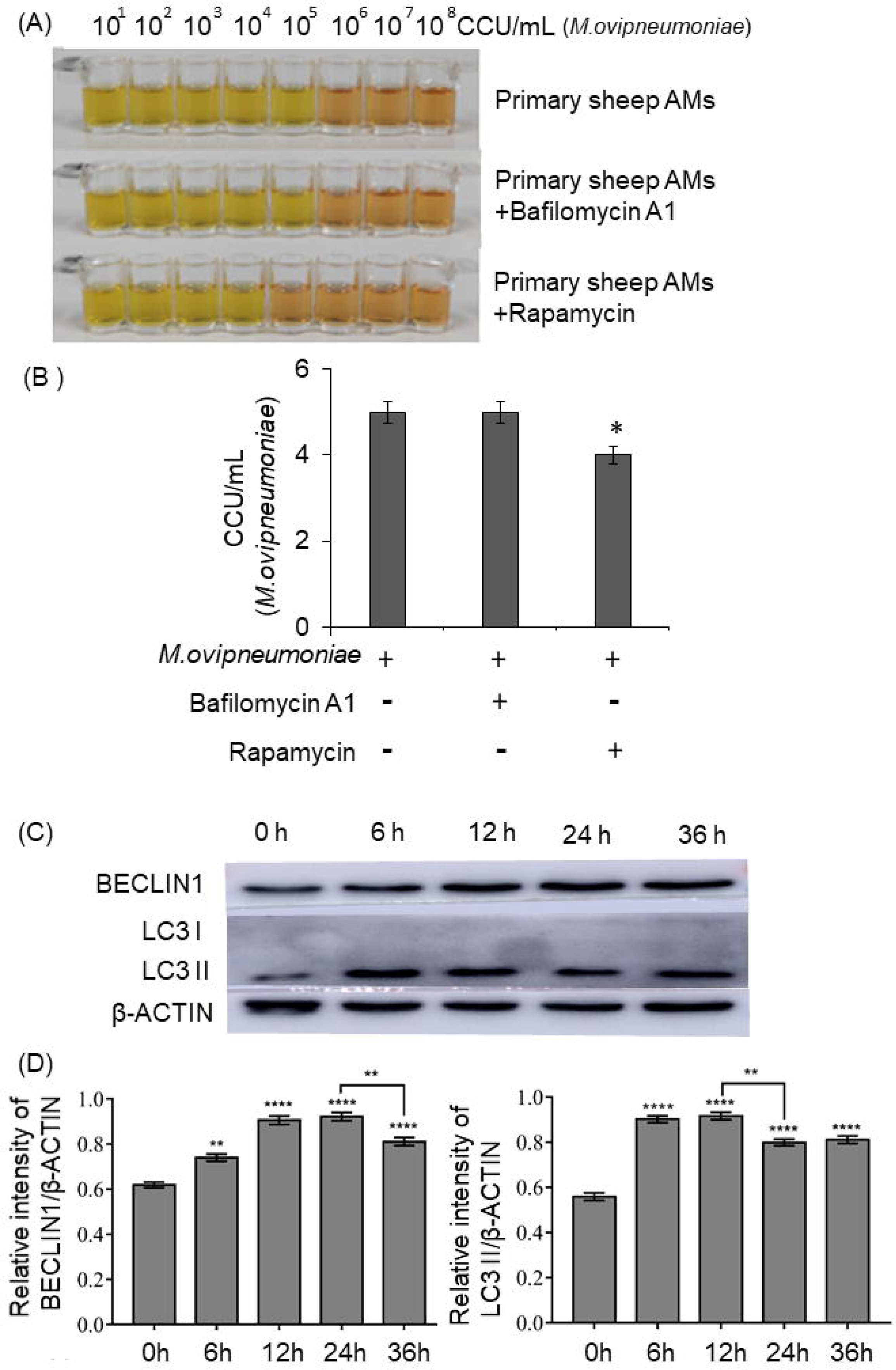

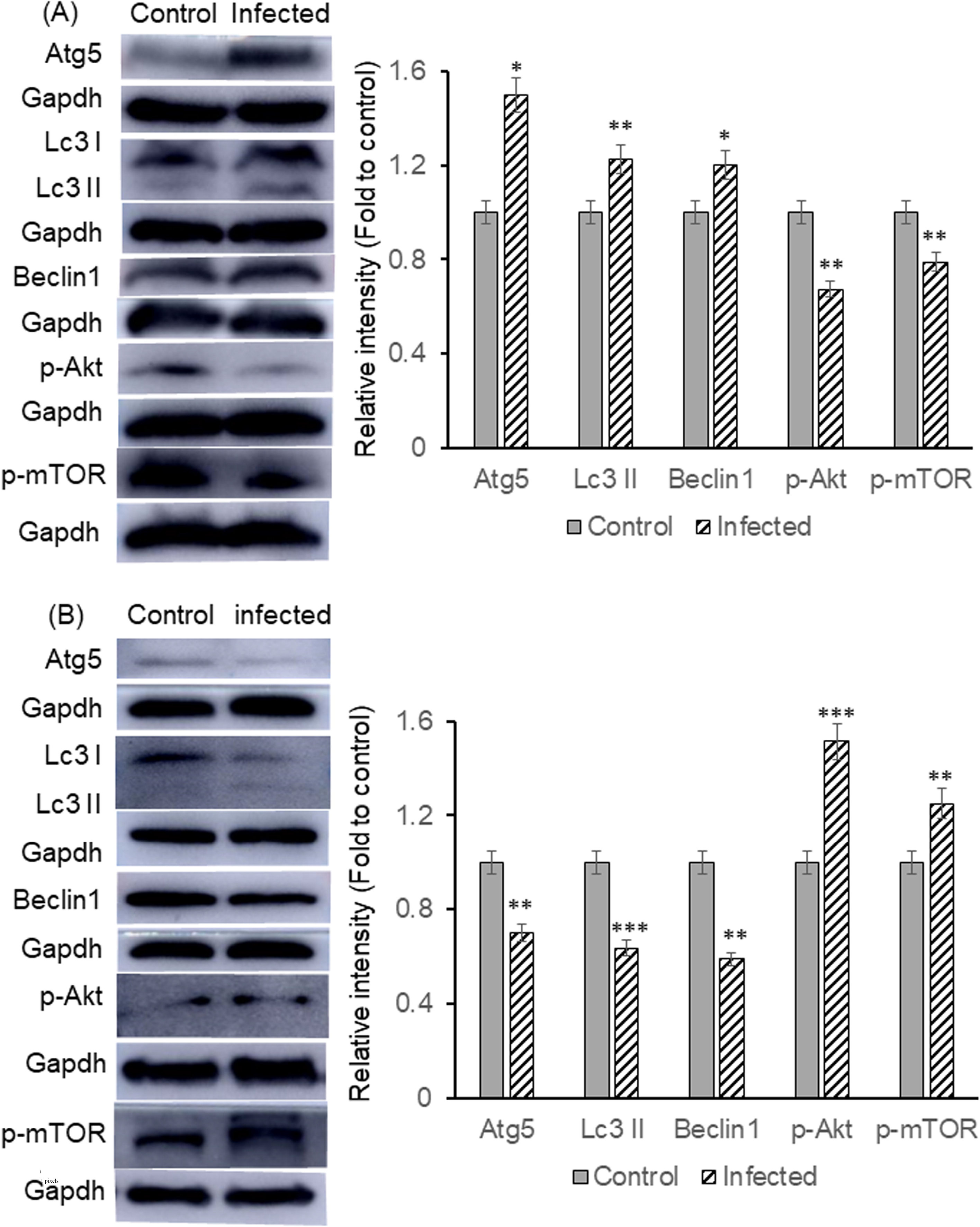

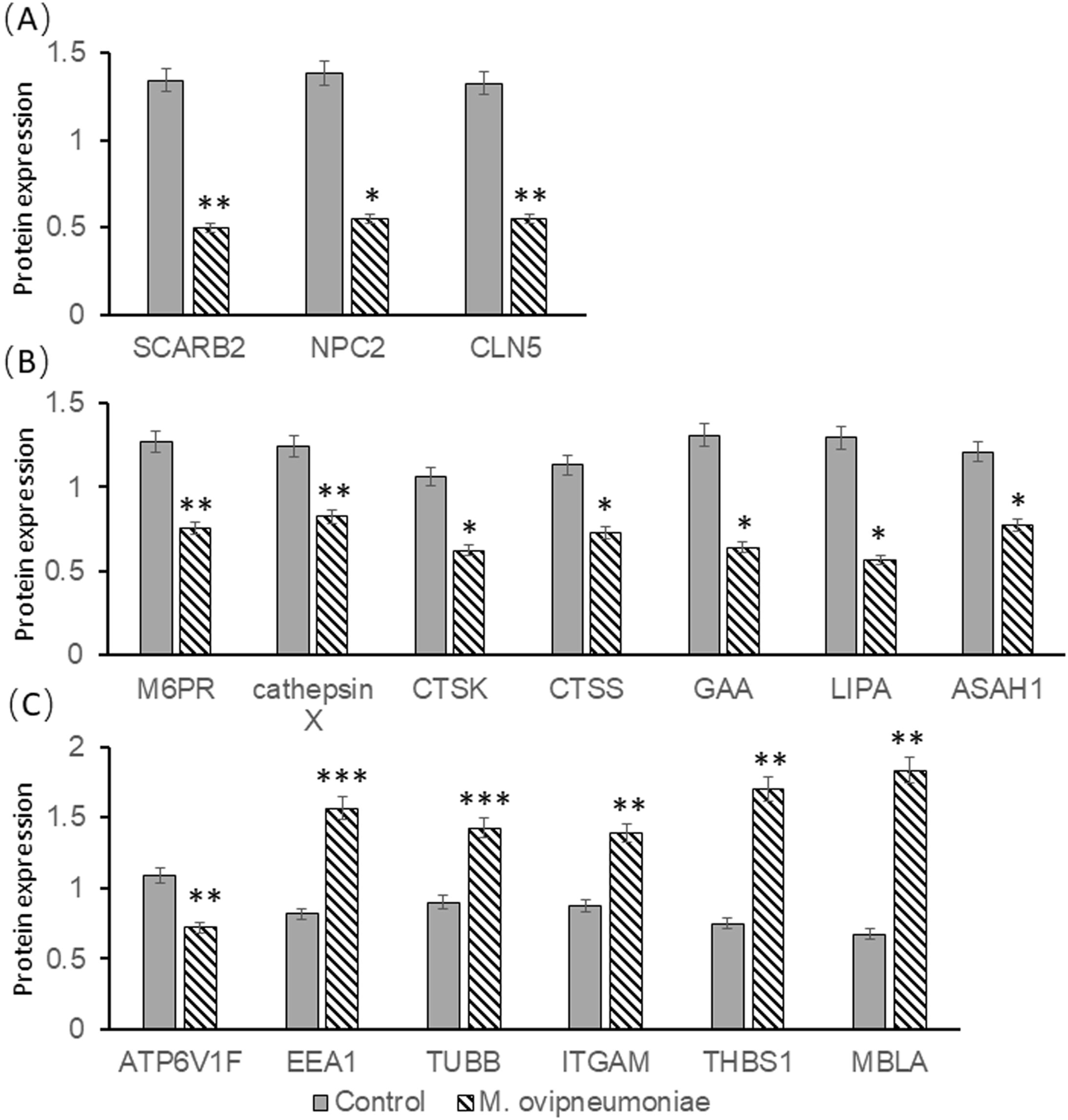

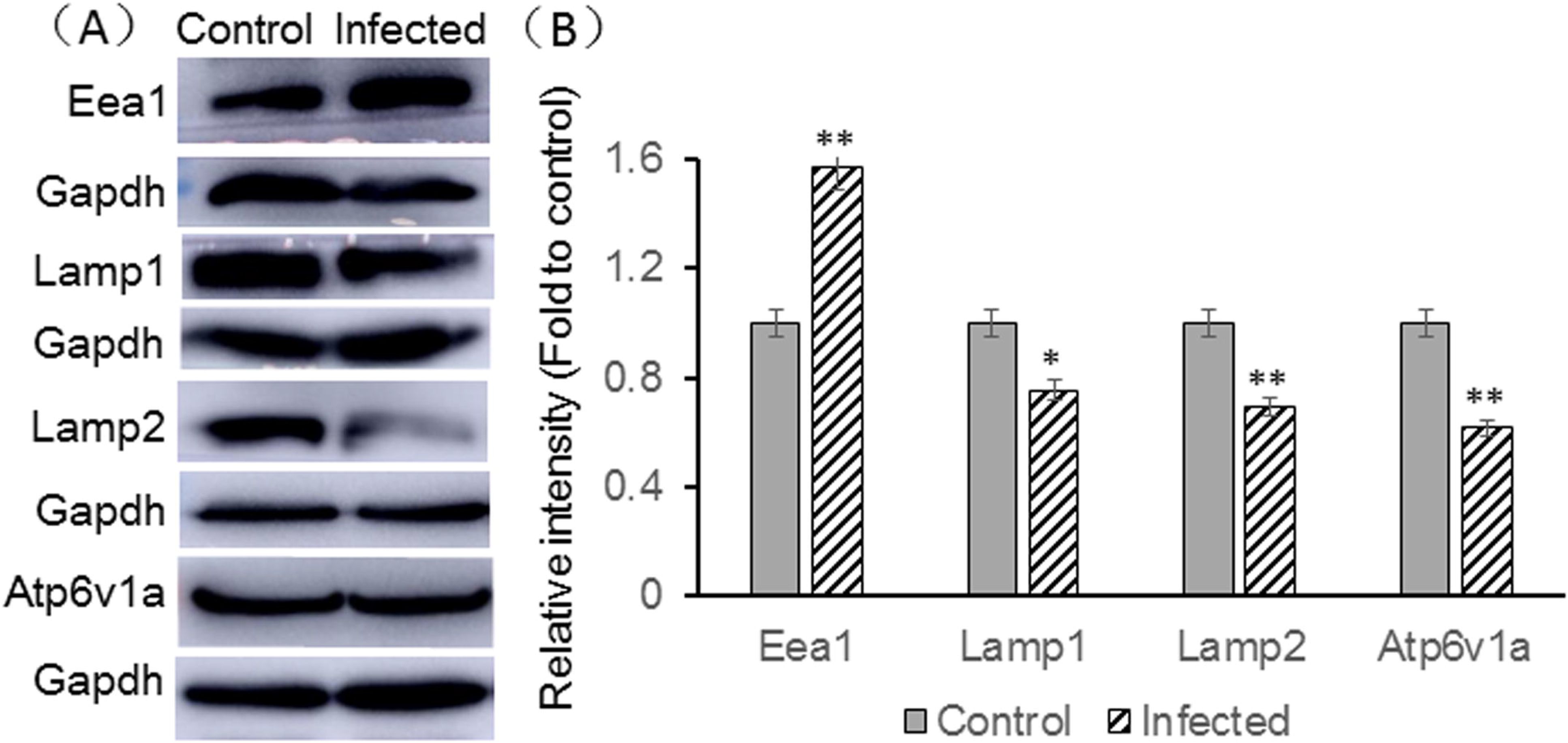

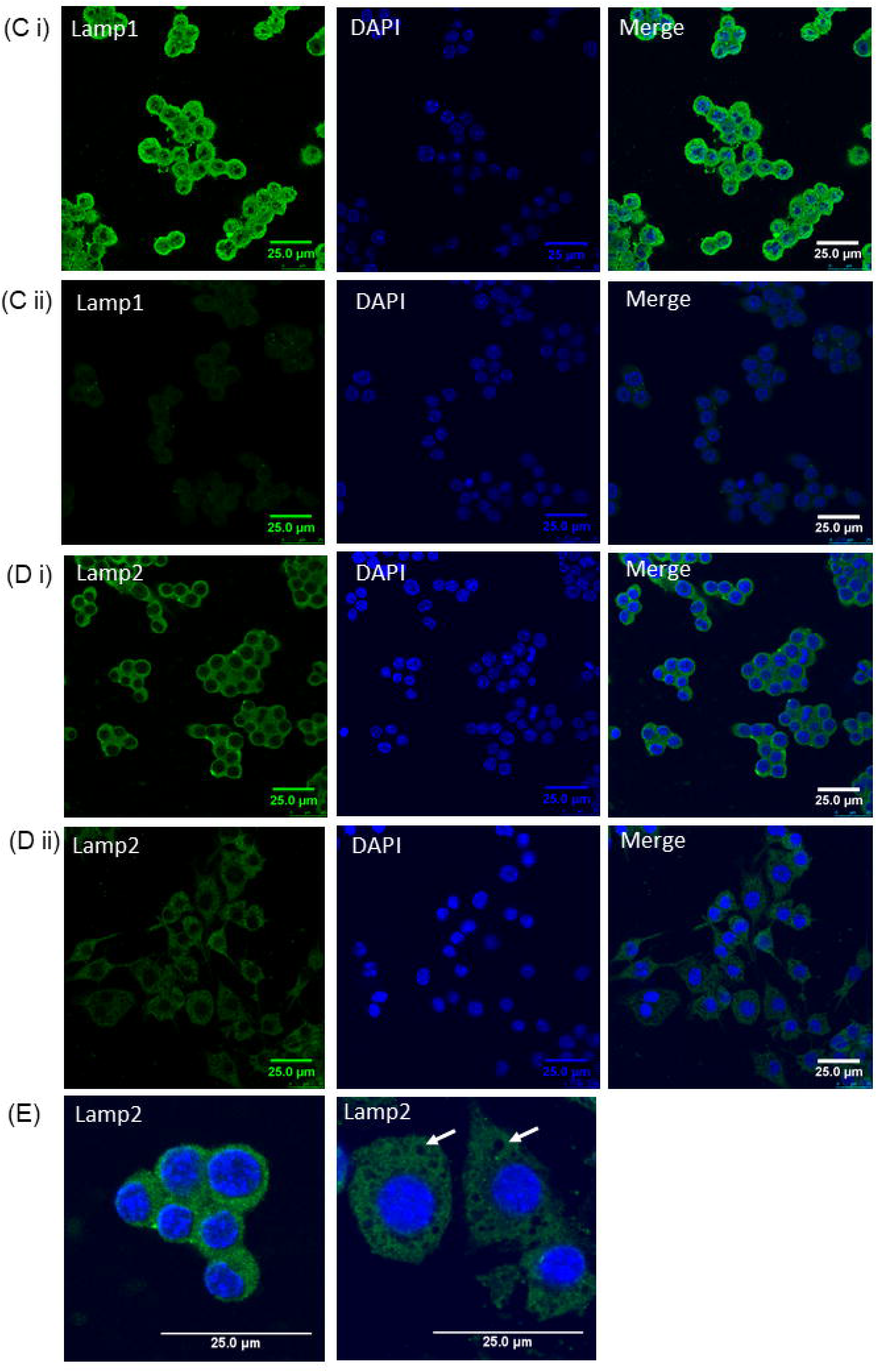

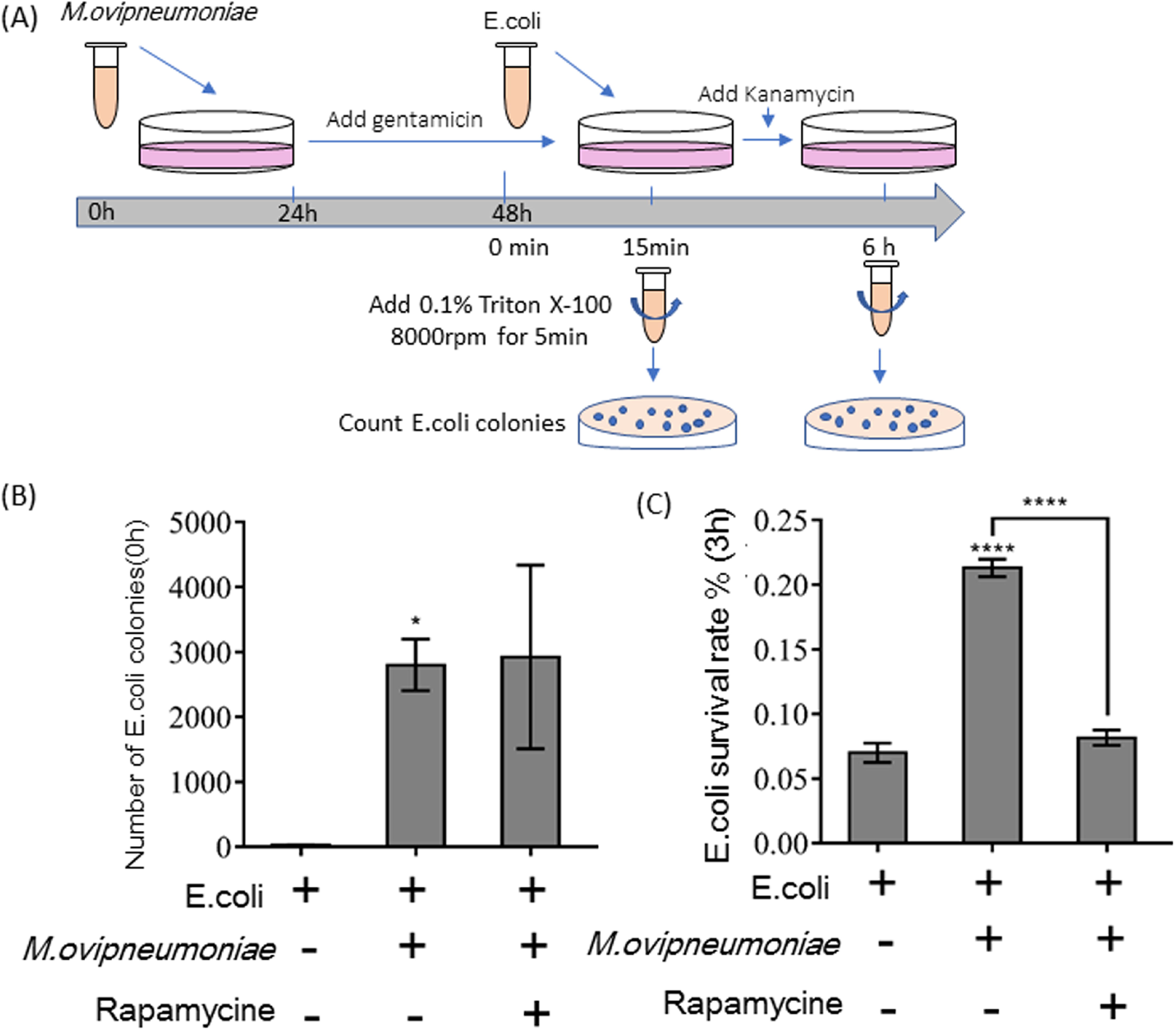

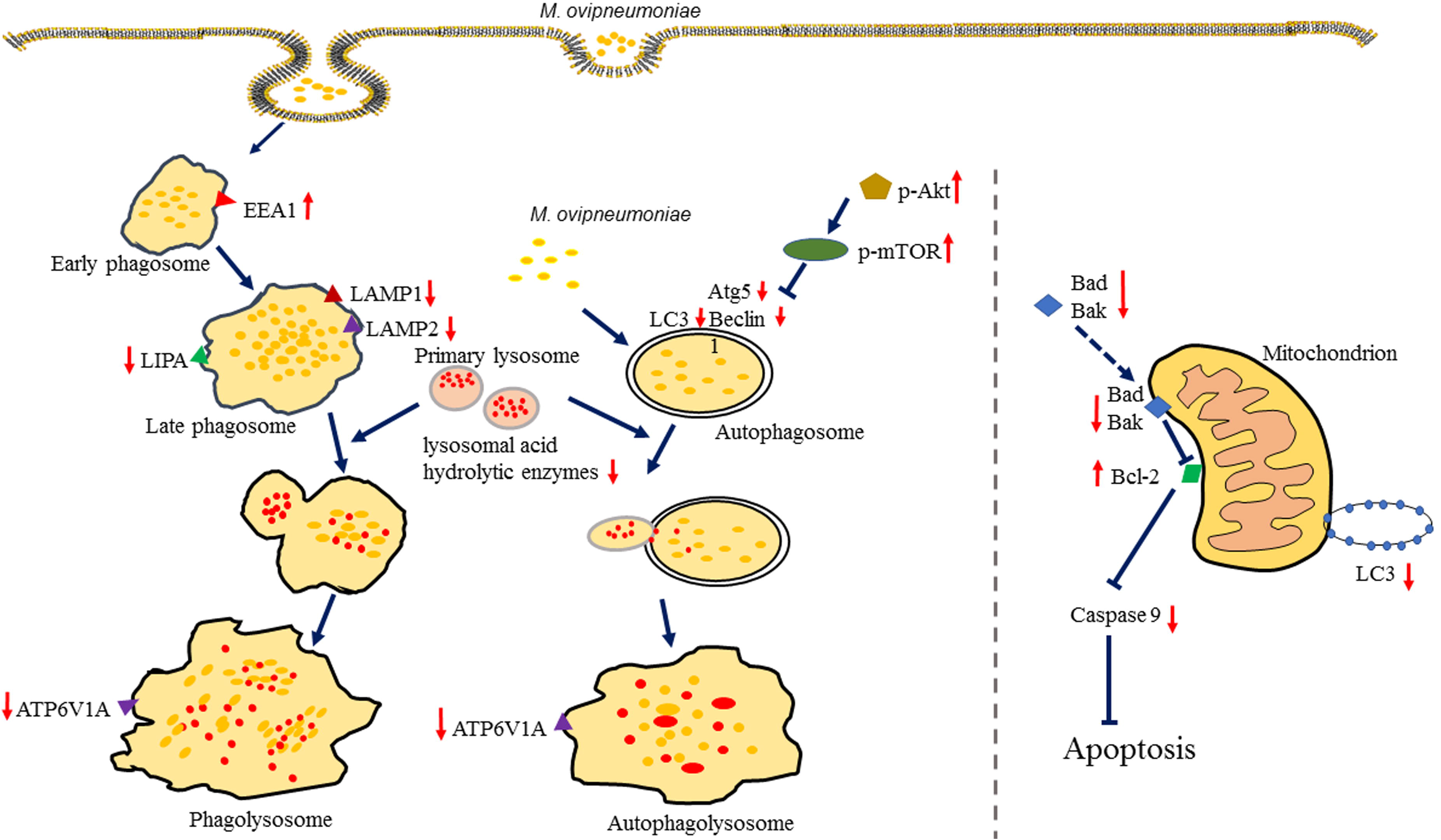

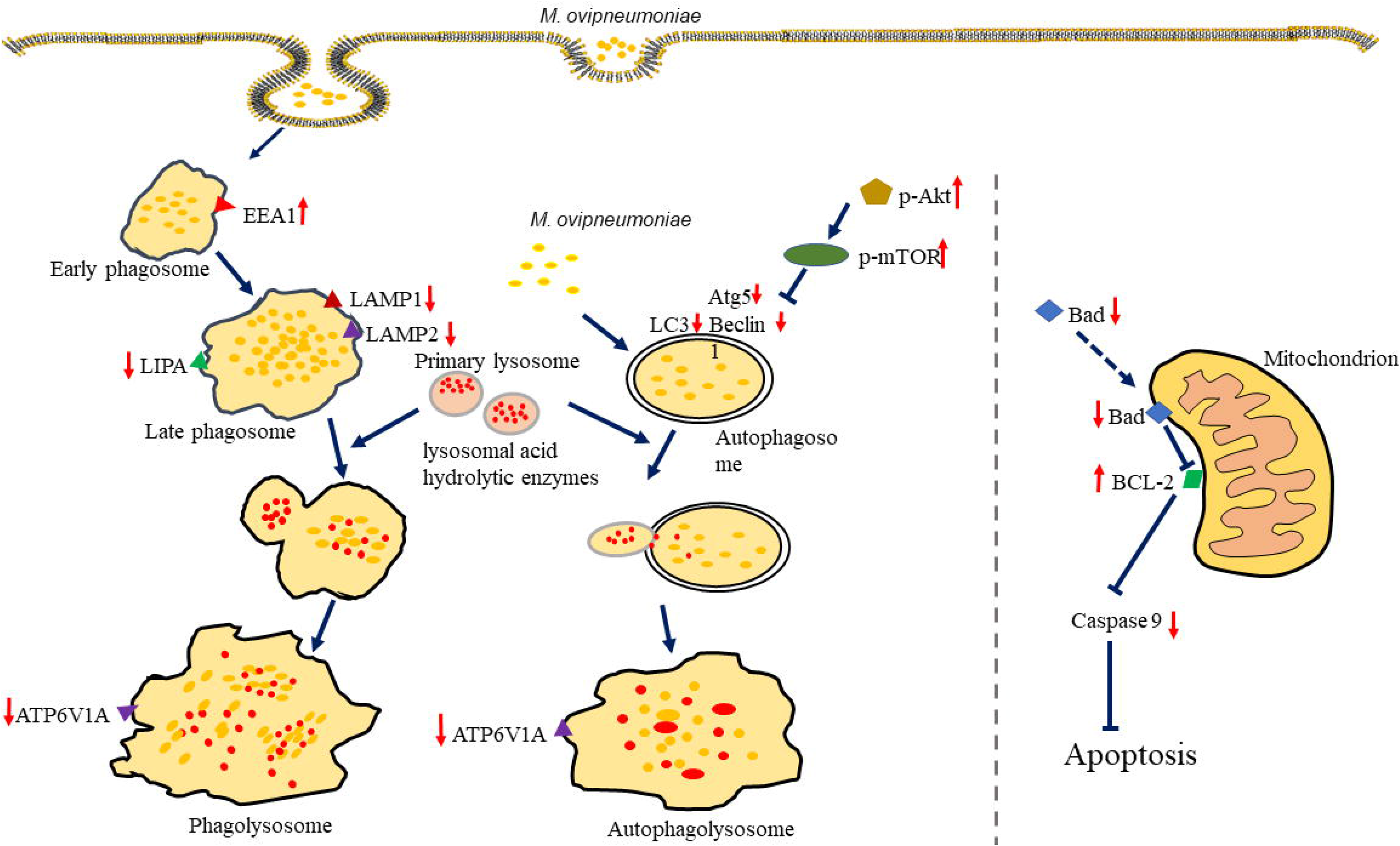

## REFERENCES

Al-Zeer, M. A., Xavier, A., Abu Lubad, M., Sigulla, J., Kessler, M., Hurwitz, R., & Meyer, T. F. (2017). Chlamydia trachomatis Prevents Apoptosis Via Activation of PDPK1-MYC and Enhanced Mitochondrial Binding of Hexokinase II. EBioMedicine, 23, 100–110.

Bento, C. F., Renna, M., Ghislat, G., Puri, C., Ashkenazi, A., Vicinanza, M., … Rubinsztein, D. C. (2016). Mammalian Autophagy: How Does It Work? Annual Review of Biochemistry, 85(1), 685–713.

Besser, T. E., Cassirer, E. F., Potter, K. A., Lahmers, K., Oaks, J. L., Shanthalingam, S., … Foreyt, W. J. (2014). Epizootic pneumonia of bighorn sheep following experimental exposure to Mycoplasma ovipneumoniae. PLoS ONE, 9(10).

Bewley, M. A., Naughton, M., Preston, J., Mitchell, A., Holmes, A., Marriott, H. M., … Dockrell, D. H. (2014). Pneumolysin activates macrophage lysosomal membrane permeabilization and executes apoptosis by distinct mechanisms without membrane pore formation. MBio, 5(5), 1–13.

Bürgi, N., Josi, C., Bürki, S., Schweizer, M., & Pilo, P. (2018). Mycoplasma bovis co-infection with bovine viral diarrhea virus in bovine macrophages. Veterinary Research, 49(1), 1–11.

Chao, M. P., Jaiswal, S., Weissman-Tsukamoto, R., Alizadeh, A. A., Gentles, A. J., Volkmer, J., … Weissman, I. L. (2010). Calreticulin is the dominant pro-phagocytic signal on multiple human cancers and is counterbalanced by CD47. Science Translational Medicine, 2(63), 63ra94.

Du, P., Li, S.-J., Ojcius, D. M., Li, K.-X., Hu, W.-L., Lin, X., … Yan, J. (2018a). A novel Fas-binding outer membrane protein and lipopolysaccharide of Leptospira interrogans induce macrophage apoptosis through the Fas/FasL-caspase-8/-3 pathway. Emerging Microbes & Infections, 7(1), 135.

Du, P., Li, S., Ojcius, D. M., Li, K., Hu, W., Lin, X., … Yan, J. (2018b). A novel Fas-binding outer membrane protein and lipopolysaccharide of Leptospira interrogans induce macrophage apoptosis through the Fas / FasL-caspase-8 / - 3 pathway. Emerging Microbes & Infections, 7(135), 1–17.

Feng, M., Chen, J. Y., Weissman-tsukamoto, R., Volkmer, J., & Yi, P. (2015). Macrophages eat cancer cells using their own calreticulin as a guide□: Roles of TLR and Btk. PANS, 112(7), 2145–2150.

Gao Li-yang, Zhang Ying, Chen Fang-zheng, Li Xiao-jie, Ma Jin-cheng, Ma Xiao-ming, Zhao Yan-long, Li M. (2020). Effect of Mycoplasma ovipneumoniae on the activity and inflammatory response of human alveolar type II epithelial cells. Chinese Journal of Preventive Veterinary Medicine, 42(10), 1051–1057.

Gilberti, R. M., & Knecht, D. A. (2014). Macrophages phagocytose nonopsonized silica particles using a unique microtubule-dependent pathway. Molecular Biology of the Cell, 26(3), 518–529.

Highland, M. A., Herndon, D. R., Bender, S. C., Hansen, L., Gerlach, R. F., & Beckmen, K. B. (2018). Mycoplasma ovipneumoniae in Wildlife Species beyond Subfamily Caprinae. Emerging Infectious Diseases, 24(12), 10–17.

Holloway, G., Fleming, F. E., & Coulson, B. S. (2018). MHC class I expression in intestinal cells is reduced by rotavirus infection and increased in bystander cells lacking rotavirus antigen. Scientific Reports, 8(1), 1–12.

Hu, B., Zhang, Y., Jia, L., Wu, H., Fan, C., Sun, Y., … Zhou, J. (2015). Binding of the pathogen receptor HSP90AA1 to avibirnavirus VP2 induces autophagy by inactivating the AKT-MTOR pathway. Autophagy, 11(3), 503–515.

Hu, W.-L., Dong, H.-Y., Li, Y., Ojcius, D. M., Li, S.-J., & Yan, J. (2017). Bid-Induced Release of AIF/EndoG from Mitochondria Causes Apoptosis of Macrophages during Infection with Leptospira interrogans. Frontiers in Cellular and Infection Microbiology, 7(November), 1–13.

Huang, H., Kang, R., Wang, J., Luo, G., Yang, W., & Zhao, Z. (2013). Hepatitis C virus inhibits AKT-tuberous sclerosis complex (TSC), the mechanistic target of rapamycin (MTOR) pathway, through endoplasmic reticulum stress to induce autophagy. Autophagy, 9(2), 175–195.

Jiang, Z., Song, F., Li, Y., Xue, D., Deng, G., Li, M., … Wang, Y. (2017). Capsular Polysaccharide is a Main Component of Mycoplasma ovipneumoniae in the Pathogen-Induced Toll-Like Receptor-Mediated Inflammatory Responses in Sheep Airway Epithelial Cells. Mediators of Inflammation, 2017, 9891673.

Jiang, Z., Song, F., Li, Y., Xue, D., Zhao, N., Zhang, J., … Wang, Y. (2017). Capsular Polysaccharide of Mycoplasma ovipneumoniae Induces Sheep Airway Epithelial Cell Apoptosis via ROS-Dependent JNK/P38 MAPK Pathways. Oxidative Medicine and Cellular Longevity, 2017, 6175841.

Kissing, S., Hermsen, C., Repnik, U., Nesset, C. K., Von Bargen, K., Griffiths, G., … Saftig, P. (2015). Vacuolar ATPase in Phagosome-Lysosome Fusion. The Journal of Biolo, 290(22), 14166–14180.

Lee, K., Whang, J., Choi, H., Son, Y., Jeon, H. S., Back, Y. W., … Kim, H. (2016). Mycobacterium avium MAV2054 protein induces macrophage apoptosis by targeting mitochondria and reduces intracellular bacterial growth. Nature Publishing Group, (November), 1–16.

Levine, B., Mizushima, N., & Virgin, H. W. (2011). Autophagy in immunity and inflammation. Nature, 469(7330), 323–335.

Li, Y., Jiang, Z., Xue, D., Deng, G., Li, M., Liu, X., & Wang, Y. (2016). Mycoplasma ovipneumoniae induces sheep airway epithelial cell apoptosis through an ERK signalling-mediated mitochondria pathway. BMC Microbiology, 16(1), 222.

Liu, X., Xu, N., & Zhang, S. (2013). Calreticulin is a microbial-binding molecule with phagocytosis-enhancing capacity. Fish and Shellfish Immunology, 35(3), 776–784.

Livingston, C. W. J., & Gauer, B. B. (1979). Isolation of Mycoplasma ovipneumoniae from Spanish and Angora goats. American Journal of Veterinary Research, 40(3), 407–408.

Martinez, J., Almendinger, J., Oberst, A., Ness, R., Dillon, C. P., Fitzgerald, P., … Green, D. R. (2011). Microtubule-associated protein 1 light chain 3 alpha (LC3)-associated phagocytosis is required for the efficient clearance of dead cells. Proceedings of the National Academy of Sciences, 108(42), 17396–17401.

Mehto, S., Antony, C., Khan, N., Arya, R., & Selvakumar, A. (2015). Mycobacterium tuberculosis and Human Immunodeficiency Virus Type 1 Cooperatively Modulate Macrophage Apoptosis via Toll Like Receptor 2 and Calcium Homeostasis, 1–20.

Mohan, K., Obwolo, M. J., & Hill, F. W. (1992). Mycoplasma ovipneumoniae infection in Zimbabwean goats and sheep. Journal of Comparative Pathology, 107(1), 73–79.

Niedergang, F., Colucci-guyon, E., Dubois, T., Raposo, G., & Chavrier, P. (2002). ADP ribosylation factor 6 is activated and controls membrane delivery during phagocytosis in macrophages. The Journal of Cell Biology, 161, 1143–1150.

Nolan, T. J., Gadsby, N. J., Hellyer, T. P., Templeton, K. E., McMulla, R., McKenna, J. P., … Simpson, A. J. (2016). Low-pathogenicity Mycoplasma spp. alter human monocyte and macrophage function and are highly prevalent among patients with ventilator-acquired pneumonia. Thorax, 71(7), 594–600.

Pimentel-Muiños, F. X., & Boada-Romero, E. (2014). Selective autophagy against membranous compartments: Canonical and unconventional purposes and mechanisms. Autophagy, 10(3), 397–407.

Purcell, R. H., Taylor-Robinson, D., Wong, D., & Chanock, R. M. (1966). Color test for the measurement of antibody to T-strain mycoplasmas. Journal of Bacteriology, 92(1), 6–12.

Rifatbegovic, M., Maksimovic, Z., & Hulaj, B. (2011). Mycoplasma ovipneumoniae associated with severe respiratory disease in goats. The Veterinary Record, 168(21), 565.

Rong, G., Zhao, J.-M., Hou, G.-Y., & Zhou, H.-L. (2014). Seroprevalence and molecular detection of Mycoplasma ovipneumoniae in goats in tropical China. Tropical Animal Health and Production, 46(8), 1491–1495.

Sanjuan, M. A., & Green, D. R. (2008). Eating for Good Health--Linking Autophagy and Phagocytosis in Host Dedense. Autophagy, 4(5), 607–611.

Tavares, R., & Pathak, S. K. (2017). Helicobacter pylori Secreted Protein HP1286 Triggers Apoptosis in Macrophages via TNF-Independent and ERK MAPK-Dependent Pathways. Frontiers in Cellular and Infection Microbiology, 7(February), 1–16.

Weiser, G. C., Drew, M. L., Cassirer, E. F., & Ward, A. C. S. (2012). Detection of Mycoplasma ovipneumoniae and M. arginini in bighorn sheep using enrichment culture coupled with genus- and species-specific polymerase chain reaction. Journal of Wildlife Diseases, 48(2), 449–453.

Xue, D., Li, Y., Jiang, Z., Deng, G., Li, M., Liu, X., & Wang, Y. (2017). A ROS-dependent and Caspase-3-mediated apoptosis in sheep bronchial epithelial cells in response to Mycoplasma Ovipneumoniae infections. Veterinary Immunology and Immunopathology, 187, 55–63.

Yang, A., Pantoom, S., & Wu, Y. (2017). Elucidation of the anti-autophagy mechanism of the Legionella effector RavZ using semisynthetic LC3 proteins. ELIFE, 6, e23905.

Yu-shin Sou, Satoshi Waguri, Jun-ichi Iwata, Takashi Ueno, Tsutomu Fujimura, T. H., Sawada, N., Yamada, A., Noboru Mizushima, Uchiyama, Y., Eiki Kominami, K. T., & Komatsu, M. (2008). Membrane Ruffles Capture C3bi-opsonized Particles in Activated Macrophages. Molecular Biology of the Cell, 19(4762–4775), 4628–4639.

