## Supplementary figures and images for "Strategies of *Mycoplasma ovipneumoniae* to evade immune clearance by alveolar macrophages"

### Supplemental Fig 1

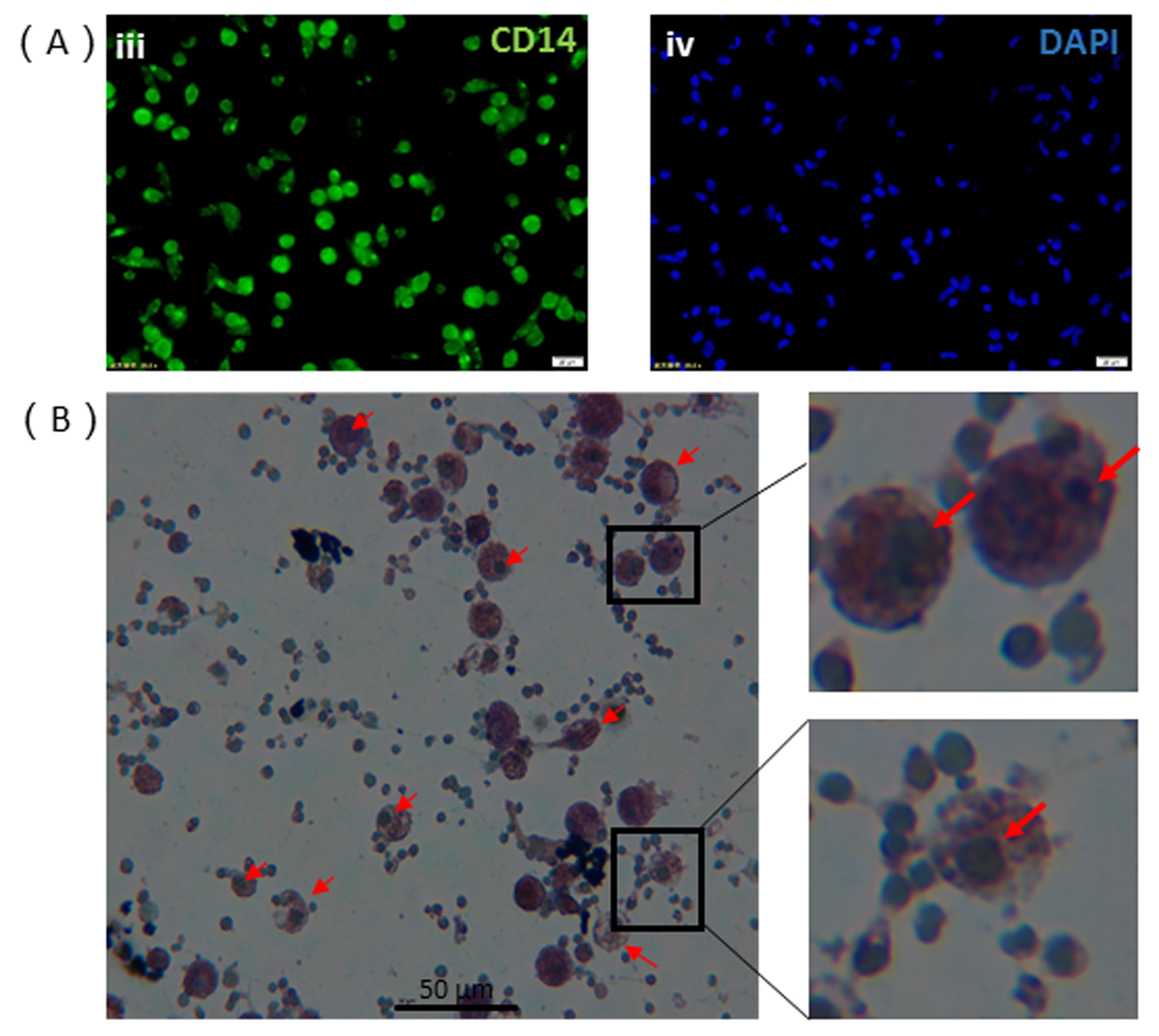

### Supplemental Fig 2

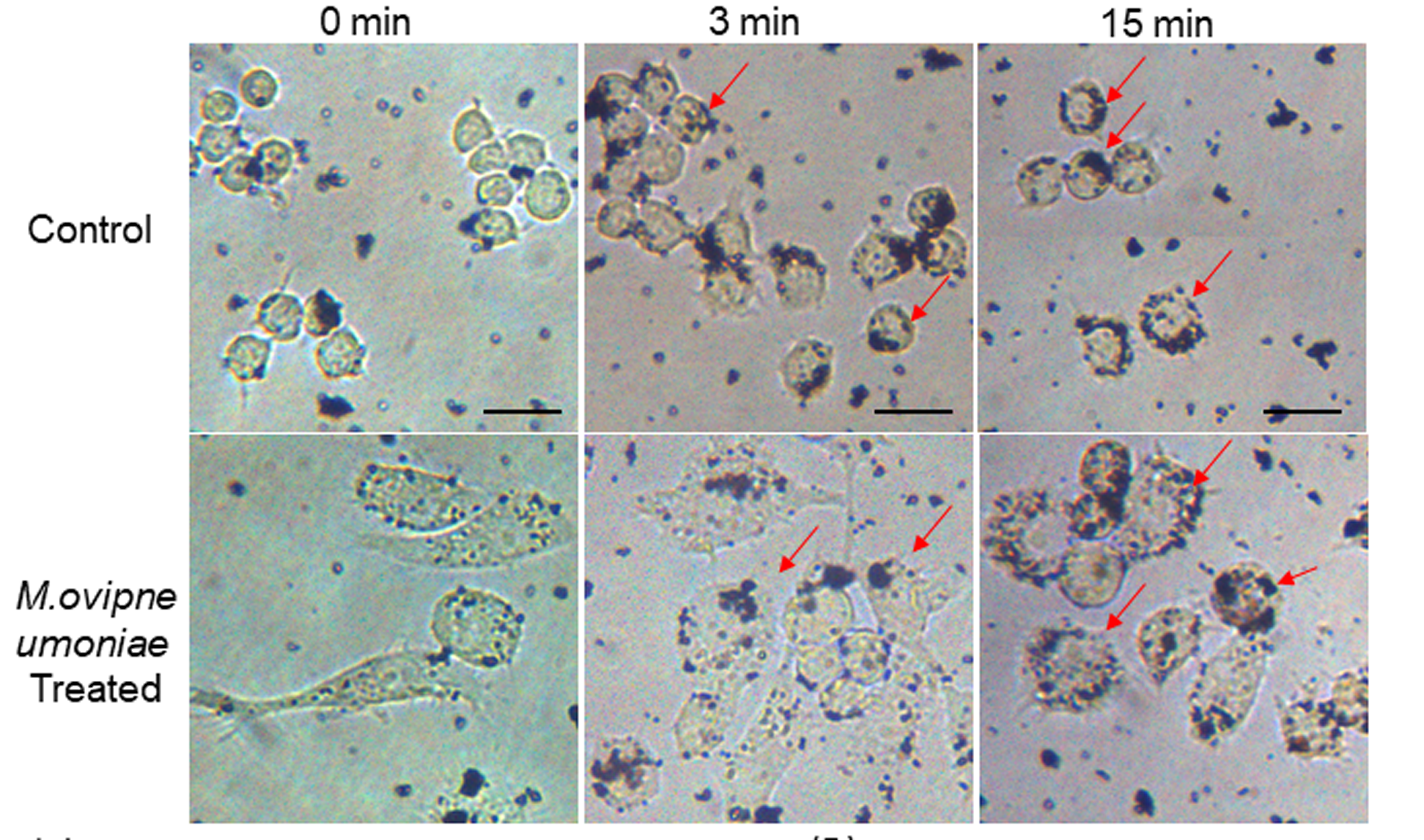

### Supplemental Fig 3

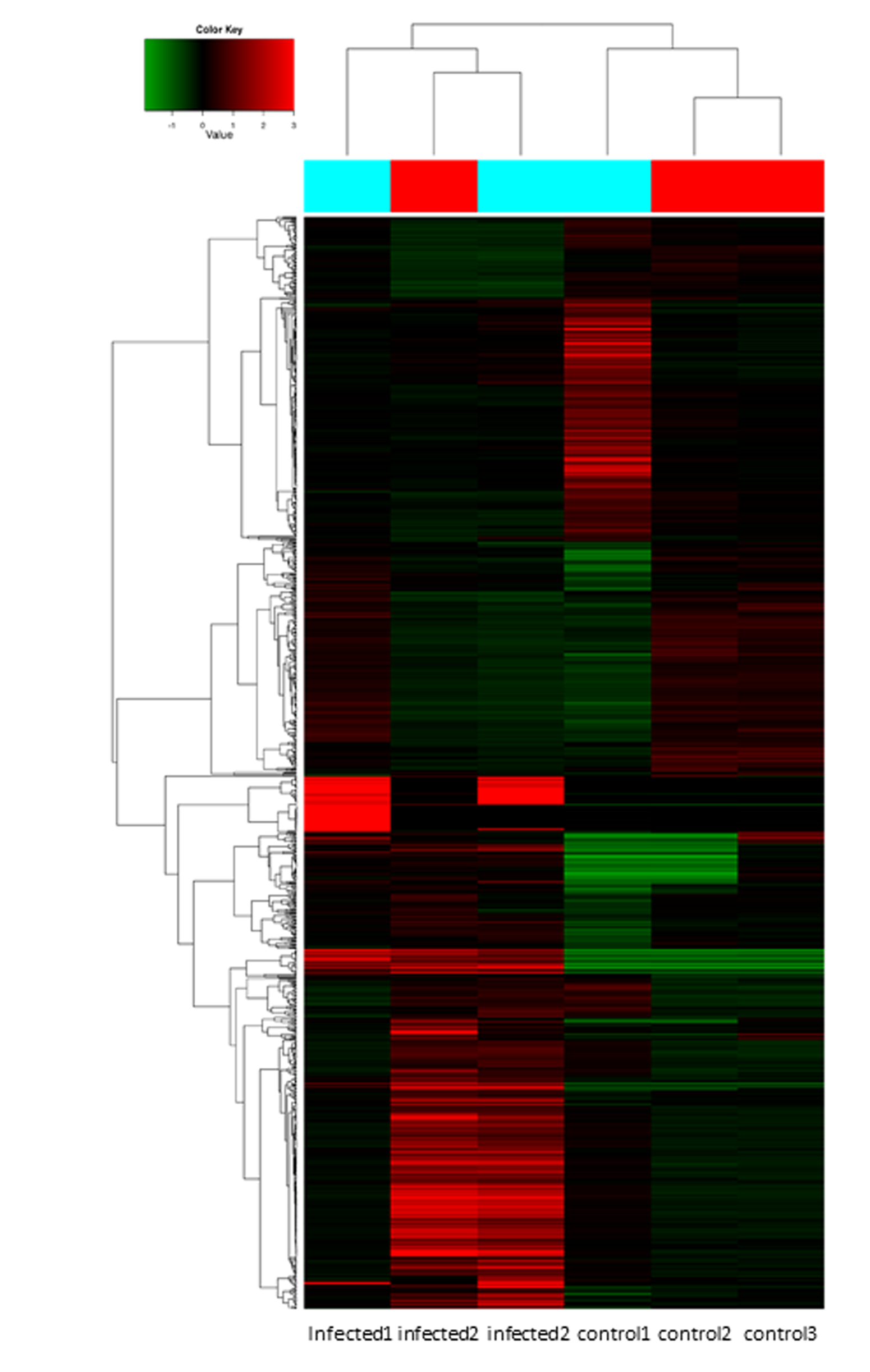

### Supplemental Fig 4

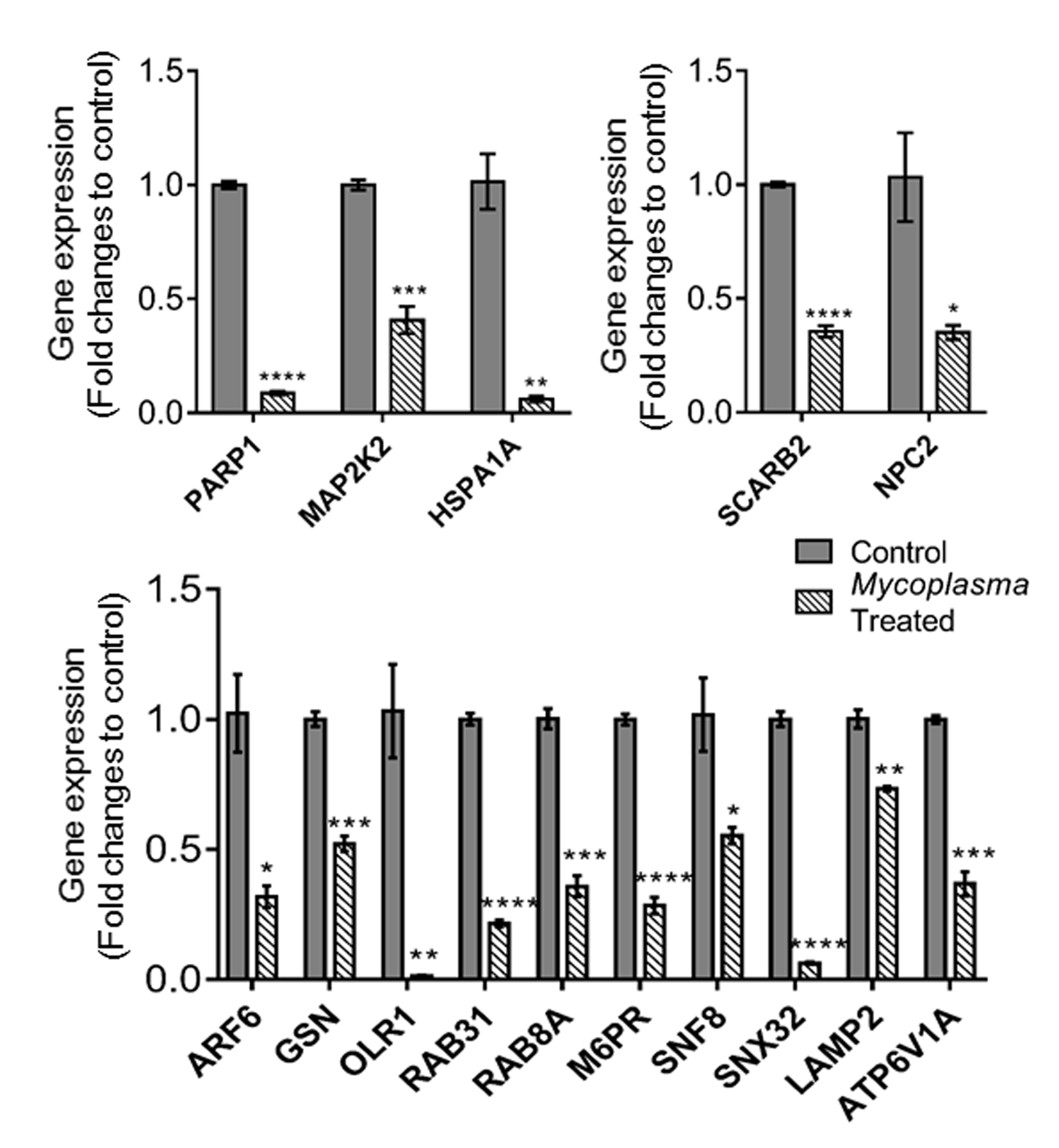

### Supplemental Fig 5

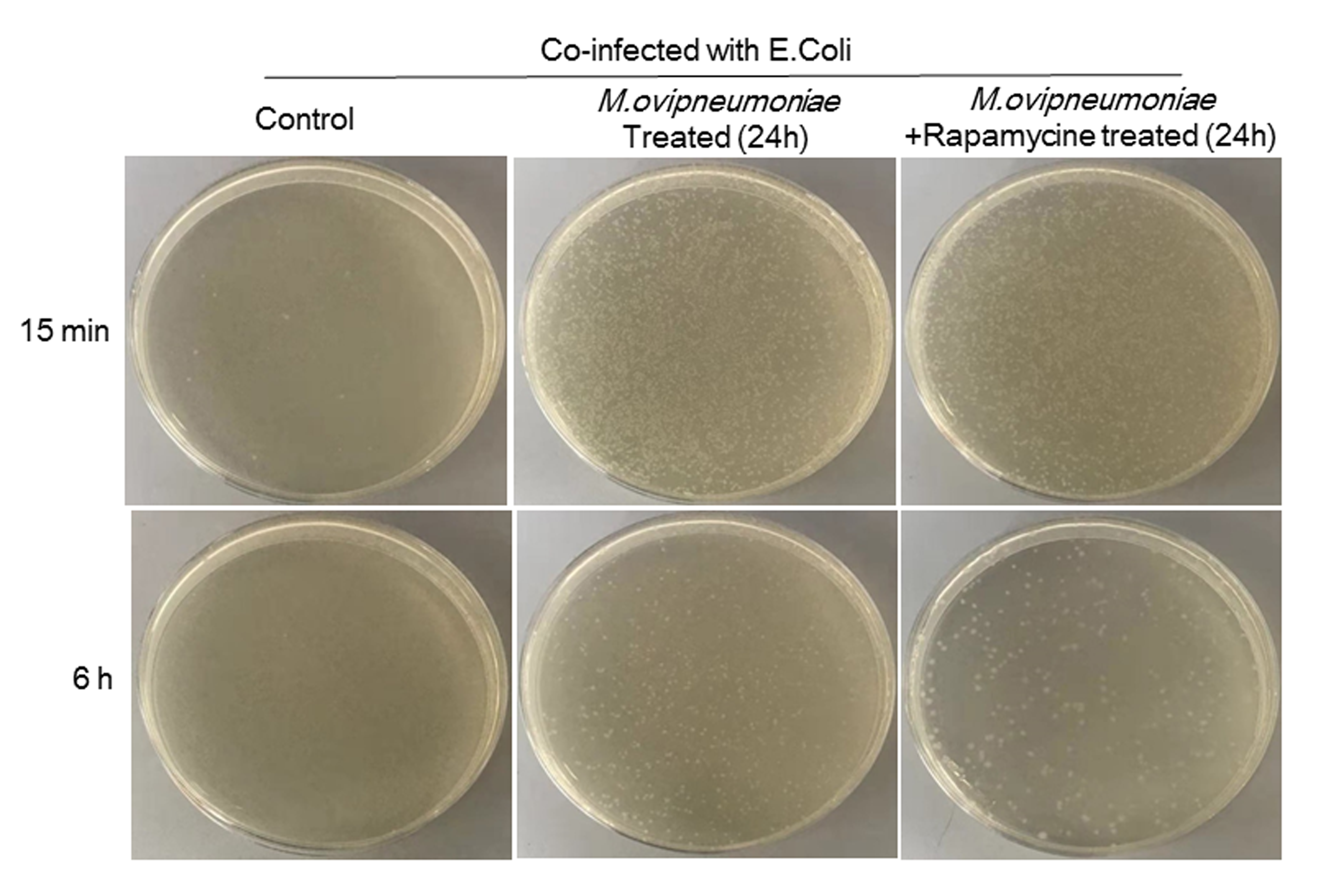
